# Single-Domain SARS-CoV-2 S1 and RBD Antibodies Isolated from Immunized Llama Effectively Bind Targets of the Wuhan, UK, and South African Strains in vitro

**DOI:** 10.1101/2021.02.15.431198

**Authors:** Divora Yemane, Ivan Lu, Winson Tiahjono, Lauren Rubidoux, Abbas Hussain, John C. Cancilla, Erika Duggan, Nathan C. Shaner, Nobuki Nakanishi, Jiwu Wang

**Affiliations:** Allele Biotechnology and Pharmaceuticals, Inc., 6404 Nancy Ridge Drive, San Diego, CA 92121; Scintillon Institute, 6868 Nancy Ridge Drive, San Diego, CA 92121

**Keywords:** COVID-SARS-2, Single Domain Antibody, Nanoantibody, Nanobody, Variants, UK Strain, South African Strain, Antigen Test, mNeonGreen

## Abstract

The spreading of SARS-CoV-2 variants has become a major challenge of the current fight against the pandemic. Of particular concerns are the strains that have arisen from the United Kingdom (UK) and South Africa. The UK variant spreads rapidly and is projected to overtake the original strain in the US as early as in March 2021, while the South African variant appears to evade some effects of the current vaccines. Potential false-negative diagnosis using currently available antigen kits that may not recognize these variants could cause another wave of community infection. Therefore, it is imperative that antibodies used in the detection kits are validated for binding against these variants. Here we report that the nanoantibodies (nAbs in our terminology, also referred to as VHH fragments, single domain antibodies, nanobodies™) that we have developed for rapid antigen detection test bind the receptor binding domain (RBD) of the S1 protein from the original COVID-SARS-2 virus as well as those from the UK and South African variants. This finding validates our antibodies used in our assay for the detection of these major variant strains.

## Main Text

We immunized a llama by repeated injections of recombinant S1 and RBD proteins. Using our nanoantibody (nAb) development pipeline that resulted in dozens of drug candidate nAbs over a decade [1], we isolated multiple families of anti-RBD nAbs that showed strong binding affinity and specificity against the RBD of the clinical strain Wuhan-Hu-1. We then characterized the properties of these nAbs through various experiments that will be published separately. We further developed a prototype rapid antigen test in the format of lateral flow assay with these nAbs using clinical samples before the spread of the current major variants.

The UK variant, also referred to as SARS-CoV-2 B.1.1.7 lineage, contains ~20 amino acid changes in total, one of which is in the RBD (N501Y) and several in the N-terminus of the S1 protein. The South Africa variant, also known as B.1.351 lineage, contains N501Y and two more changes in RBD (K417N, E484K). These alterations might impact the epitope binding of conventional monoclonal antibody (mAb) pairs selected for existing antigen detection tests. Nanoantibodies, on the other hand, bind their antigens through distinct mechanism than IgG antibodies. With a size that is approximately 10x smaller than IgG, the VHH single-domain derived nAbs can insert its epitope-contacting amino acids into crevices on their antigens [2] instead of “wrapping around” exposed epitopes by a heavy chain/light chain arm pair of IgG, affording a possibility that the epitope recognition of nAbs might be less affected by individual amino-acid mutations of the variants than that of mAbs. In this report, we show that our nAbs indeed bind RBDs of both the UK and the South Africa lineages.

Recombinant RBDs of Wuhan-Hu-1 (WT), B.1.1.7 (UK), and B.1.351 (South Africa) were obtained from BEI and conjugated to Spherotech particles before flow cytometry was performed. We tested the binding ability of nAb1 and nAb2, representing two of several families of anti-SARS-CoV-2 nAbs that we isolated. We previously determined their dissociation constant (Kd) toward WT RBD through serial dilutions as a measure of binding affinity and found that nAb1 and nAb2 showed Kds between 6 nM to 15nM when conjugated to different fluorescent proteins. We then set to test the binding affinities of nAb1 and nAb2 towards variant RBDs as described below. To facilitate the detection of nAbs in the flowcytometry assay as well as our rapid antigen diagnostic kit, we genetically fused each nAb with mNeonGreen, the brightest monomeric fluorescent protein that we previously developed [3]. We also produced a genetic fusion protein that contains two nAb1 domains linked via a linker (referred to as nAb1 dimer).

As shown in Figure 1, we found that both mNeonGreen-fused nAb1 and nAb2 bind to RBDs from all three strains above the background binding to bovine serum albumin (BSA). Note that these three RBDs when conjugated to the Spherotech beads as carrier particles have not been normalized for quantitative cross-comparison, therefore we could not conclude relative binding affinity for each RBD. More quantitative and direct comparison will be performed with additional experiments. As a control, we used the monoclonal antibody CR3022, which was originally isolated for its binding to the S protein of SARS-CoV [4] and crossreacts to SARS-CoV-2 S protein. CR3022 we obtained from R&D Systems binds to the WT RBD but not to UK or South African RBDs in our hand, despite the computational prediction that mutations on South African RBD might minimally affect CR3022 binding [5]. Interestingly but not surprisingly, nAb1 dimer binds all RBDs with affinity seemingly higher (after adjusting different fluorescent labelling) than the nAb1 monomer, indicating that the dimerization of nAb1 produces cooperative binding perhaps to RBD proteins juxtaposed on the same bead. Since individual SARS-CoV-2 virions contain 24+9 S protein trimers [6], such cooperative binding by the nAb dimer (and potentially trimer, data not shown) would likely increase the detection sensitivity of virus particles in antigen test kit. Incidentally, the benefit of creating nAb1 linear multimers have been independently demonstrated in their increased ability to neutralize SARS-CoV-2 infection using our human iPSC-derived lung epithelial cell infection experiments, to be published elsewhere.

**Figure 1.**
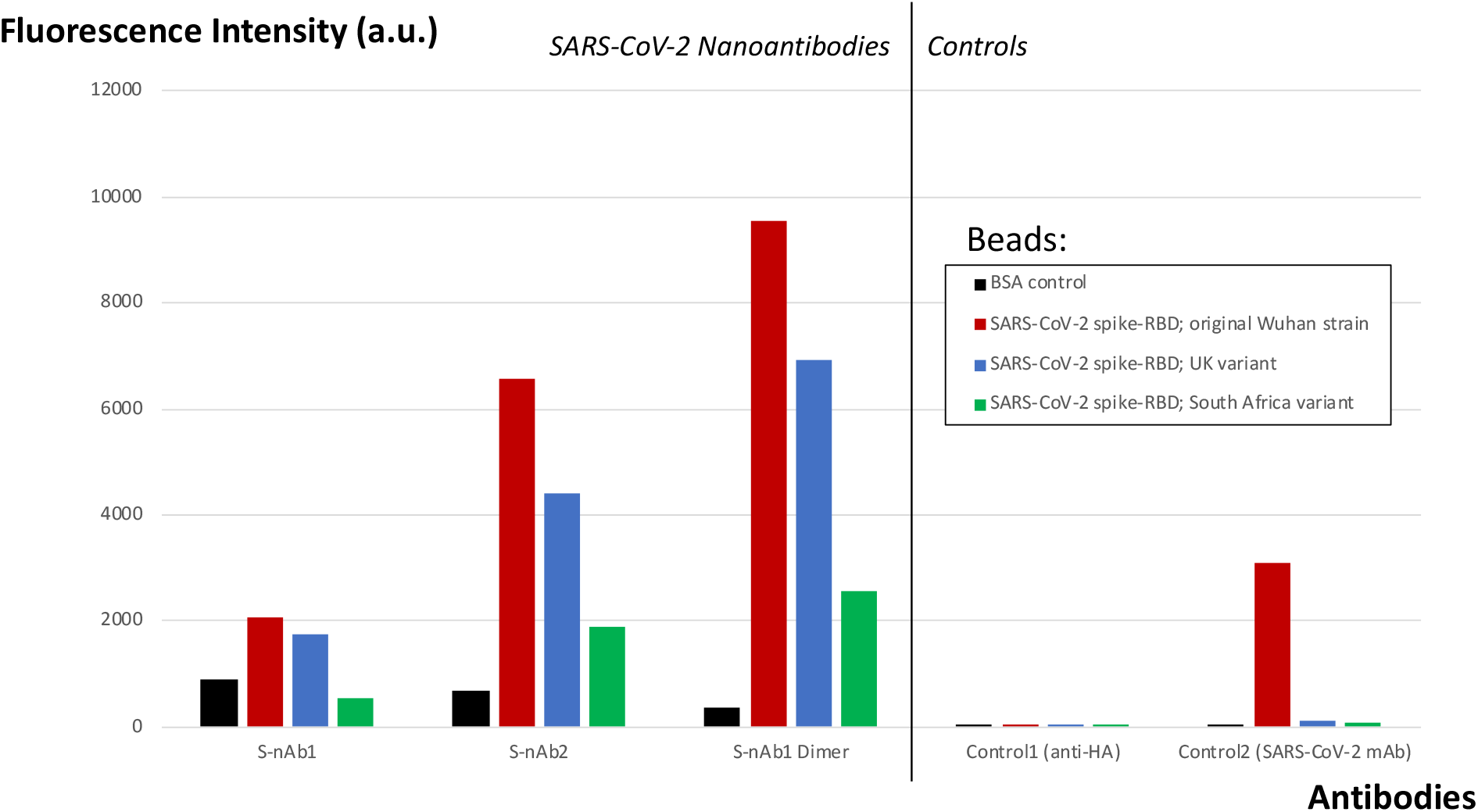
Graphical representation of binding assay results obtained via flow cytometry. We evaluated the ability of nAbs to bind spike Receptor Binding Proteins (RBPs) from SARS-CoV-2 Wuhan, UK and South African strains via. The results presented in the left panel indicate that S-nAb1 binds to Wuhan and UK RBPs, but not to South African RBP; while S-nAb2 and S-nAb1 Dimer bind to all three (Wuhan, UK, South Africa) RBPs. The right panel of the figure shows two control experiments. In control 1 (negative control), the anti-HA mAb alone resulted in low or no binding to BSA- or RBP- conjugated beads (within a range of 5 to 28 a.u.). In ctonrol 2, the commercially avaialbe SARS-CoV-2 mAb CR3022 binds to Wuhan RBD, but not to UK or South African RBPs.

Ongoing testing of at least three more families of nAbs selected from immunized llama should yield additional nAbs that are potential detectors of not only the two variants tested in this study but also other variants in circulation or will arise in the future. Finally, Allele Biotech’s COVID antigen rapid test, designed for both point-of-care clinical and over-the-counter home use, integrates multiple nAbs’ capabilities of recognizing hundreds of binding pockets of the spike protein on the surface of virus particles, enabling detection of current and future variants and much improved sensitivity over the existing test kits.

## Methods

### nAb production

The nAbs that bind to SARS-CoV-2 Wuhan RBD/S1 proteins were isolated by methods that we described in detail [1]. Briefly, a llama was repeatedly immnunized with recombinant Wuhan RBD and S1 proteins obtained from BEI Resources. Several weeks after the initial injection, the blood from the llama was harvested and mRNAs isolated from an enriched lymphocyte population. Reverse transcriptase (RT)-PCR was performed using a set of primers that specifically amplify VHH domains and amplified cDNAs cloned into an M13 vector. Phage display based on enzyme-linked immunosorbent assay (ELISA) was performed in 96-well plates to isolate VHH cDNAs that specificially bind to Wuhan RBD and S1 proteins. The positivie cDNAs are subcloned into an E. coli expression vector for the production of RBD/S1 nAbs.

### Flow cytometry using protein-conjugated beads

Bovine serum albumin (BSA) and three recombinant RBDs (Wuhan, UK, and South African) were covalently conjugated to Speherotech bead particles and flow cytometry was performed after the beads were mixed with three fluorescent labeled nAbs that we generated; S-nAb1, S-nAb2 and S-nAb1 Dimer. For fluorescent labeling, S-nAb1 and S-nAb2 are genetically fused with mNeonGreen [3]. S-nAb1 Dimer is HA-tagged and detected through anti-HA mAb (Cat# IC6875G, Lot# AFFY0119021, Alexa Fluor 488, Novus Biologicals). In two control experiments (1 and 2), the secondary antibody anti-HA mAb alone and the commercially available anti-SARS-CoV-2 Spike mAb (CR3022, CAT# NBP2-90980F, Lot#T2022B04-110920-F, FITC, R&D Biosystems) were tested for their ability to bind the four beads described above. Fluorescence intensity (a.u.) was adjusted per labeling methods and chromophores used in each experiment.

## Acknowledgment

The following reagents were obtained through BEI Resources, NIAID, NIH: Spike Glycoprotein S1 Domain from SARS-Related Coronavirus 2, Wuhan-Hu-1 with C-Terminal Histidine Tag, Recombinant from HEK293 Cells, NR-53798; Spike Glycoprotein Receptor Binding Domain (RBD) from SARS-Related Coronavirus 2, Wuhan-Hu-1 with C-Terminal Histidine Tag, Recombinant from HEK293 Cells, NR-53800; Spike Glycoprotein Receptor Binding Domain (RBD) from SARS-Related Coronavirus 2, United Kingdom Variant with C-Terminal Histidine Tag, Recombinant from HEK293 Cells, NR-54004; Spike Glycoprotein Receptor Binding Domain (RBD) from SARS-Related Coronavirus 2, South Africa Variant with C-Terminal Histidine Tag, Recombinant from HEK293 Cells, NR-54005.

## Author Contributions Statement

DY, IL, WT, LR carried out the experiments; JCC designed the solubilized version of the nAbs and nAb multimers; NCS, NN and JW designed and together with AH supervised the nAb generation experiments; ED conducted and supervised flowcytometry experiment; JW wrote the main manuscript; JCC prepared the figure; NN and JW edited the draft; JW secured and provided funding.

## Data Availability

Additional, related data of the present report is available from the corresponding author upon request.

## References

1. Wong, A, Sykora, C, Rogers, L, Higginbotham, J, Wang J. Modified Nanoantibodies Increase Sensitivity in Avidin-Biotin Immunohistochemistry. 2018 Appl Immunohistochem Mol Morphol. 26, 682–688. PMID: 28059871

2. Jovcevska I & Muyidermans S. The therapeutic potential of nanobodies. 2020 BioDrugs 34, 11–26 PMID: 31686399

3. Shaner NC, Lambert GG, Chammas A, Ni Y, Cranfill PJ, et al., Wang J. A bright monomeric green fluorescent protein derived from Branchiostoma lanceolatum. 2013 Nat Methods. 10, 407–9. PMID: 23524392

4. Ter Meulen J, van den Brink EN, Poon LLM, Marissen WE, et al., Human monoclonal antibody combination against SARS coronavirus: synergy and coverage of escape mutants. 2006 PLOS Med. 3, e237 pmed.003023

5. Cheng MH, Krieger JM, Kaynak B, Arditi M, Bahar I. Impact of South African 501.V2 variant on SARS-CoV-2 spike infectivity and neutralization. A structure-based computational assessment. 2021 bioRxiv https://doi.org/10.1101/2021.01.10.426143 doi:

6. Ke Z, Oton J, Qu K, Cortese M, et al., Structures and distributions of SARS-CoV-2 spike proteins on intact virions 2020 Nature 588, 498–502

